# Immunodietica: interrogating the role of diet in autoimmune disease

**DOI:** 10.1101/2020.05.05.079418

**Authors:** Iosif M. Gershteyn, Andrey A. Burov, Brenda Y. Miao, Vasco H. Morais, Leonardo M. R. Ferreira

## Abstract

Diet is an environmental factor in autoimmune disorders, where the immune system erroneously destroys one’s own tissues. Yet, interactions between diet and autoimmunity remain largely unexplored, particularly the impact of immunogenetics, one’s human leukocyte antigen (HLA) allele make-up, in this interplay. Here, we interrogated animals and plants for the presence of epitopes implicated in human autoimmune diseases. We mapped autoimmune epitope distribution across organisms and determined their tissue expression pattern. Interestingly, diet-derived epitopes implicated in a disease were more likely to bind to HLA alleles associated with that disease than to protective alleles, with visible differences between organisms with similar autoimmune epitope content. We then analyzed an individual’s HLA haplotype, generating a personalized heatmap of potential dietary autoimmune triggers. Our work uncovered differences in autoimmunogenic potential across food sources and revealed differential binding of diet-derived epitopes to autoimmune disease-associated HLA alleles, shedding light on the impact of diet on autoimmunity.

## Main

Food has the potential to affect every aspect of one’s health. It provides not just energy and building blocks to our cells, but also many molecules with pharmacological properties, leading many to see food as medicine. Unsurprisingly, there is a push towards dissecting the composition of food at the molecular level in greater detail^1^.

The role of diet in the prevalence and progression of autoimmune disorders is poorly understood and presently understudied. There are almost 100 described autoimmune disorders, affecting an estimated 50 million Americans. Still, the etiology of most cases remains unknown. Twin studies across different countries have revealed that genetics alone can only predict 22% of cases of common autoimmune diseases^2^, leading to an increased recognition of the importance of environmental factors^3^.

Diet likely plays a key role among environmental factors affecting autoimmune disease incidence and severity. Milk consumption positively correlates with multiple sclerosis (MS) incidence across 27 countries^4^. Fasting, followed by one year of vegetarian diet, significantly reduces pain, morning stiffness, swollen joints, and other symptoms in rheumatoid arthritis (RA) patients^5^. Food hypersensitivity was detected in 39% of infants tested in double-blind placebo-controlled food challenge (DBPCFC) studies^6^. Several ways in which diet impacts autoimmunity have been defined at the molecular level. High salt intake enhances the differentiation of pathogenic Th17 cells, a subset of CD4^+^ T helper cells of the adaptive immune system involved in autoimmunity, via the SGK1 kinase, and aggravates neural inflammation^7,8^. Red meat contains a sugar absent in humans, Neu5Gc, which upon absorption in the intestine and incorporation in proteins and lipids, is recognized by xenoautoantibodies, leading to chronic inflammation^9^. Yet, the connection between the diet of autoimmune disease patients and their symptoms remains opaque.

Consumption of food-derived proteins containing autoimmune epitopes may contribute to the commonly noticed links between certain foods and autoimmune disease flares. Molecular mimicry is a phenomenon that occurs when a foreign antigen shares an immunogenic sequence, an epitope, with self-antigens^10^. Epitopes can be either recognized directly by antibodies, either soluble or on the surface of B cells, or presented by specialized molecules, human leukocyte antigens (HLA), to T cells. Within the HLA antigen presentation system, HLA class I molecules (HLA-A, -B, -C) present to cytotoxic CD8^+^ T cells, whereas HLA class II molecules (HLA-DR, -DP, -DQ) present to CD4^+^ T helper cells^11^. Instances of molecular mimicry as the cause of autoimmune disease have been described both in mouse and in human studies. Bacterial and viral infections were first implicated in the pathogenesis of human autoimmune disease for Guillain-Barré syndrome, a neurological autoimmune disease^12^. Progression of myocarditis, an autoimmune disease where T cells recognizing myosin heavy chain destroy cardiac tissue, requires priming of these cells by peptides derived from *Bacteroides* species in the intestine in mice^13^. Increased levels of high affinity food-specific IgG antibodies have been detected in the intestine and serum of RA patients^14^. Fifteen years ago, pork abattoir workers processing pig brains using compressed air developed an autoimmune neurologic disorder. Studies concluded that aerosolized pig neural antigens, of identity yet to be determined, induced polyradiculoneuropathy, a normally slow-developing disease, in 4 weeks^15^.

How are diet-derived epitopes recognized by the immune system? Most food absorption takes place in the small intestine. Intestinal barrier integrity is compromised by alcohol, processed food consumption, inflammation, and aging^16-18^. This can lead to a “leaky gut”, allowing for the release of bacterial products, such as flagellin and endotoxin, into the blood stream and concomitantly partly undigested food antigens structurally similar to self-antigens in an inflammatory context^19,20^. Blood brain barrier integrity is also compromised with aging^21^, so antibodies against diet-derived epitopes mimicking neural epitopes can reach the central nervous system (CNS) and cause damage^22^.

Here, we tackled the challenge of studying potential interactions between diet and autoimmunity by creating a comprehensive database of overlap between human autoimmune epitopes and food epitopes, mapping their tissue expression pattern, and analyzing their presentation by human leukocyte antigen (HLA) alleles associated with autoimmune disorders.

## Results

We built a database with all linear peptide epitopes implicated in human autoimmune diseases^23^ (www.iedb.org). The diseases with most identified epitopes were, in order of decreasing number of epitopes, multiple sclerosis (MS), rheumatoid arthritis (RA), celiac disease, systemic lupus erythematosus (SLE), type 1 diabetes (T1D, also known as insulin-dependent diabetes mellitus), and Behcet syndrome (Extended Data Figure 1). As expected, these same diseases also had the highest number of identified protein antigens, with the exception of celiac disease (Extended Data Figure 2, Figure 1a). Interestingly, of all autoimmune epitopes, less than 80% could be traced back to human, with part of them being only found in cereals, bacteria or viruses (Figure 1b). To determine the autoimmunogenicity of different foods, we mapped all 10,605 human autoimmune epitopes onto the proteomes of 24 organisms. Importantly, only exact matches were considered, as the degree of peptide sequence similarity does not predict molecular mimicry or cross-reactivity^24^. Organisms commonly consumed as food fall in three categories according to their autoimmune epitope content: red meat, poultry and fish, and cereals and vegetables. Curiously, the non-primate organism with the highest autoimmune epitope content was the bat (Figure 1c). Closing in at the individual disease level, we found that this pattern holds true for diseases with most known epitopes, such as MS, SLE, and RA (Figure 1d). Nonetheless, exceptions exist in diseases with less known epitopes. We divided diseases in three categories according to the number of implicated epitopes: “hot diseases” with more than 500 epitopes), warm diseases with between 50 and 500 epitopes, and cold diseases with less than 50 epitopes (Extended Data Figure 3). Inspecting diseases in each one of these categories revealed, for instance, that pig contains the most epitopes implicated in autoimmune gastritis, whereas the same is true for rabbit and demyelinating polyneuropathy, and rice for pemphigus gestationis (Figure 2).

**Figure 1.**
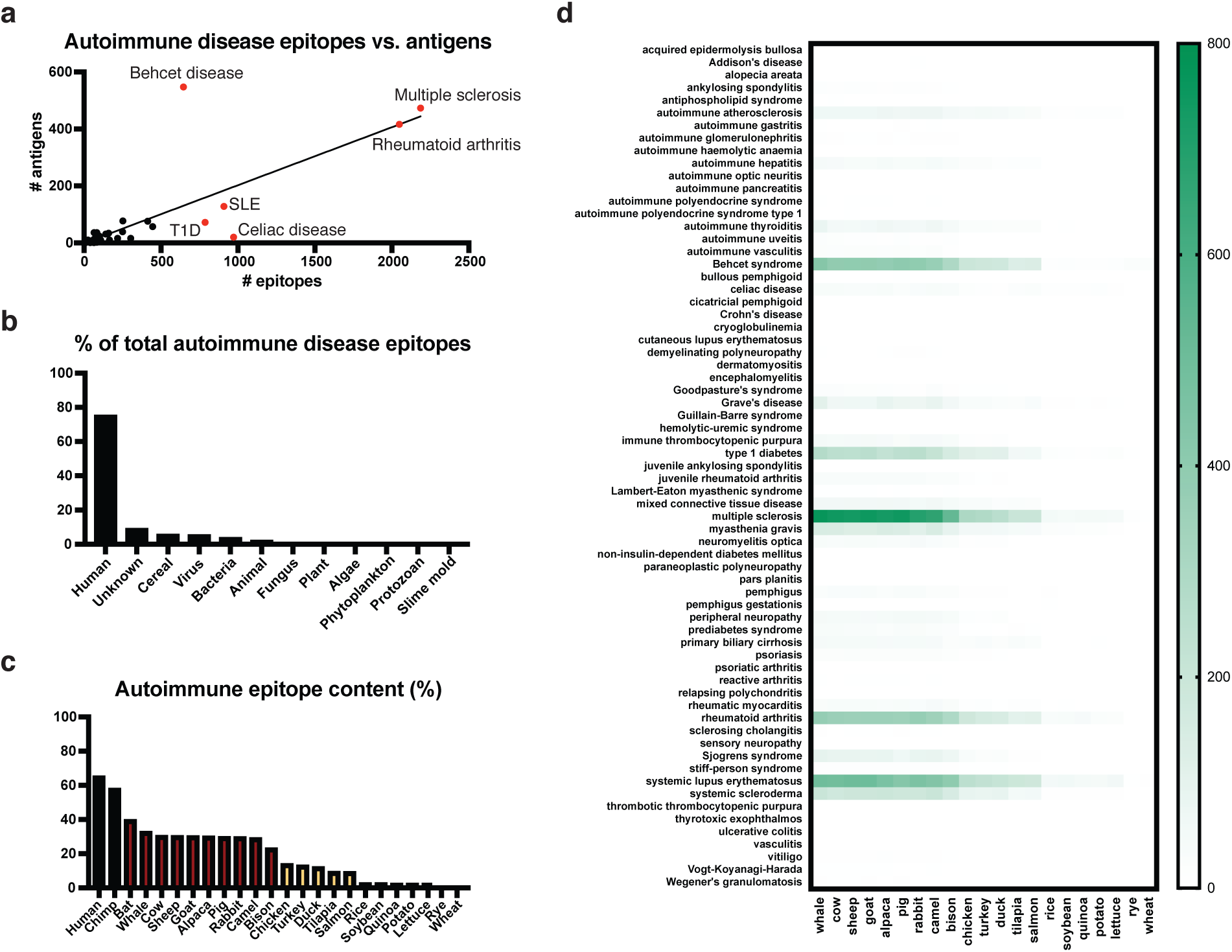
Mapping diet-derived epitopes implicated in human autoimmune disease. **a**. Correlation between the number of autoimmune epitopes and antigens across different diseases (R^2^ = 0.6620). **b**. Attributed origin of epitopes implicated in human autoimmunity (www.iedb.org). **c**. Human autoimmune epitope content of 24 species. **d**. Heat map of autoimmune epitopes present in organisms commonly consumed as food (x axis) per disease (y axis). T1D, type 1 diabetes; SLE, systemic lupus erythematosus.

**Figure 2.**
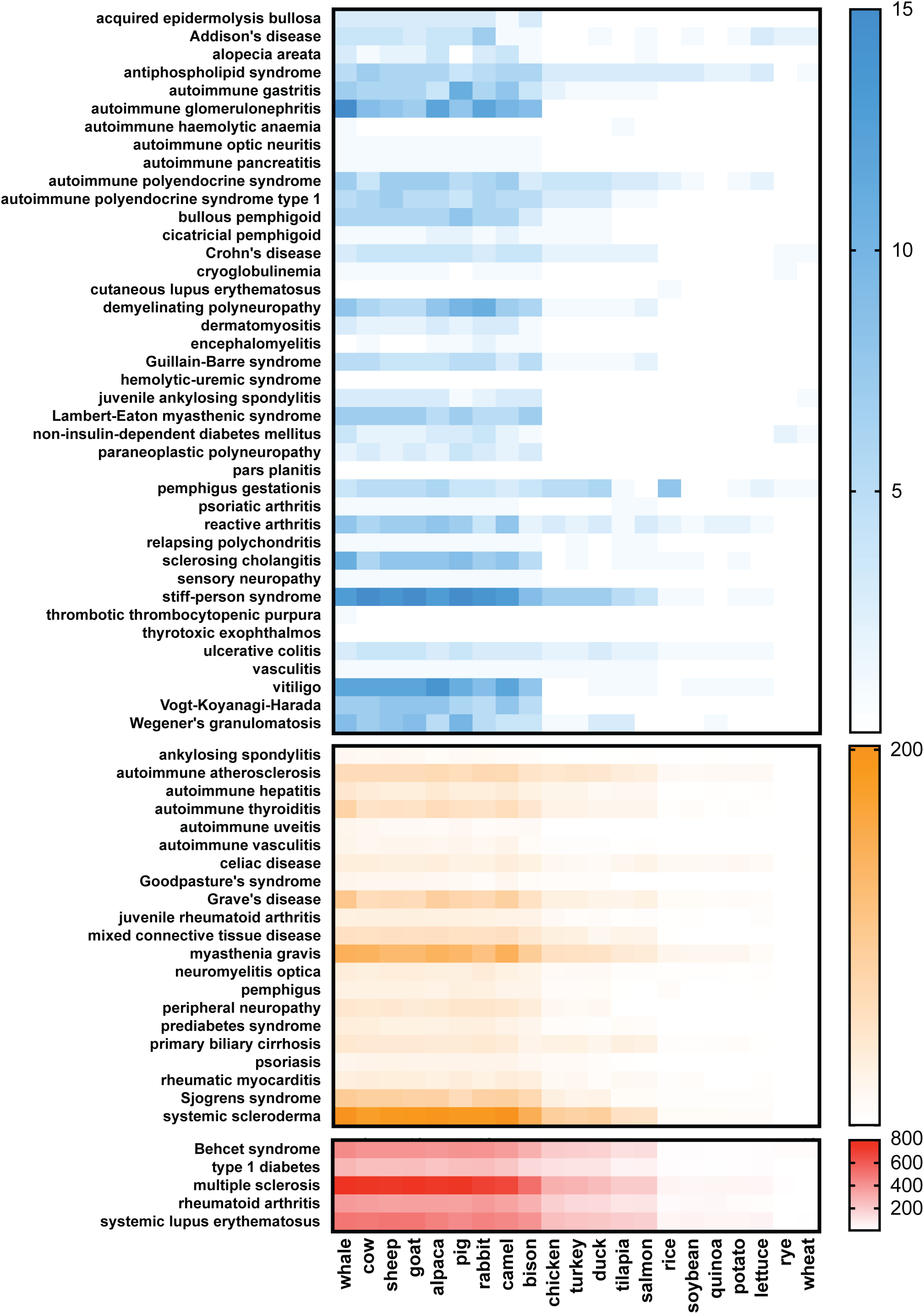
Autoimmune epitope content in food sources by disease. Three groups of autoimmune diseases (on the y axis) created based on the number of known epitopes are displayed, with diet-derived disease-specific epitopes quantified per organism (on the x axis). Organisms are aligned in order of decreasing autoimmune epitope content. Red, “hot” (> 500 epitopes); orange, “warm” (50-500 epitopes); blue, “cold” (< 50 epitopes).

Diet-derived antigens have been found to mimic a variety of human tissue-specific antigens^25^. Hence, we analyzed the expression pattern of all human autoimmune epitopes (Figure 3a), as well as of those implicated in individual diseases (Figure 3b-f). Most epitopes for MS are expressed in the brain (Figure 3b) and most of those implicated in T1D are in the pancreas (Figure 3e) can be seen. Of note, epitopes for human autoimmune disease in general are most commonly expressed in the brain (Figure 3a). Indeed, the disease with most identified epitopes is a neurological disorder, multiple sclerosis (Figure 3b, Extended Data Figure 1) and brain is also the second most common tissue of expression for T1D (Figure 3e). Strikingly, all analyzed organisms contain epitopes that map back to human proteins present in every organ, from the cow and pig to rice (Figure 4a-f).

**Figure 3.**
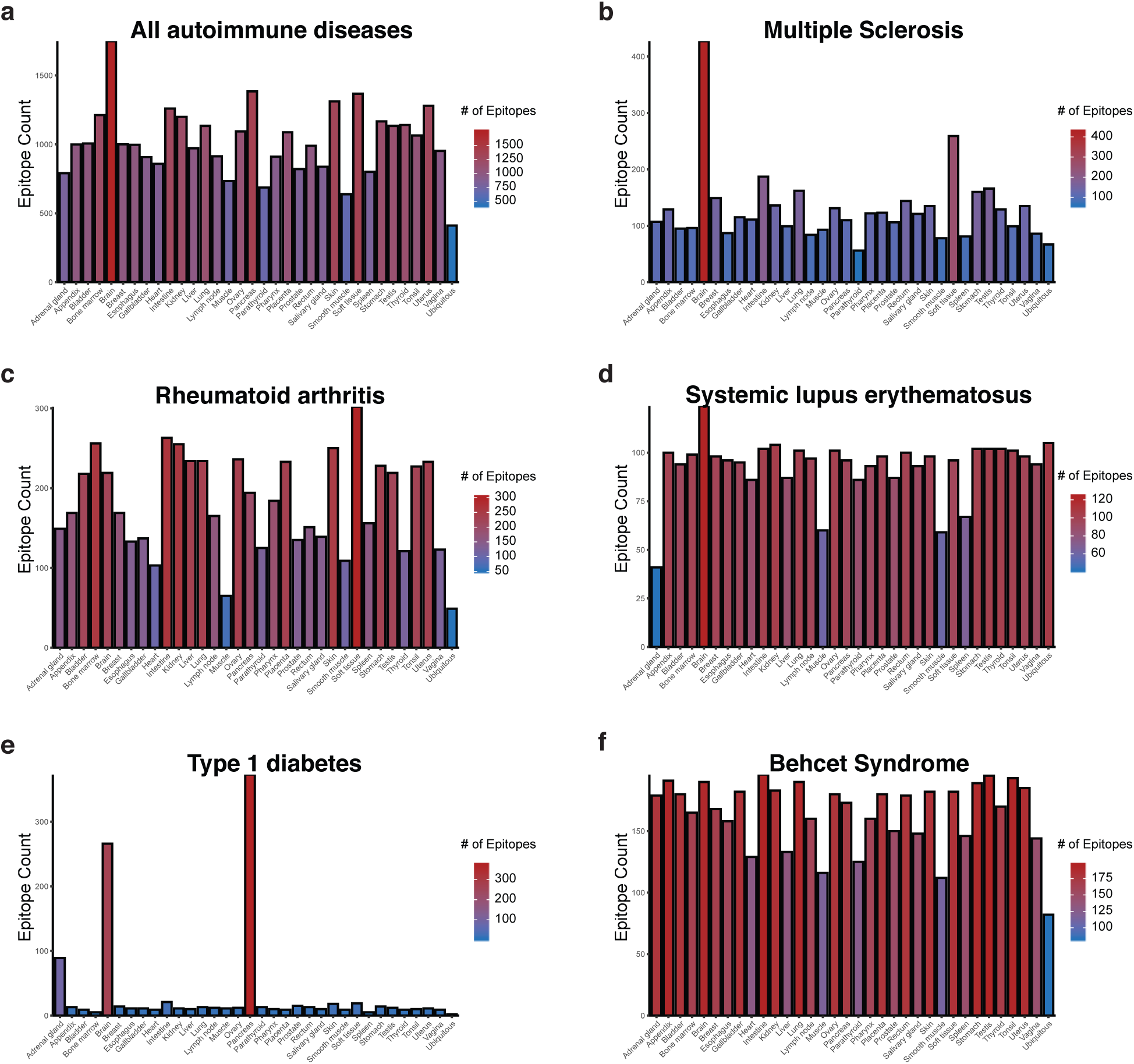
Tissue expression pattern of human autoimmune epitopes. Expression by organ of epitopes implicated in (**a**) human autoimmune disease, (**b**) multiple sclerosis, (**c**) rheumatoid arthritis, (**d**) systemic lupus erythematosus, (**e**) type 1 diabetes, (**f**) Behcet syndrome.

**Figure 4.**
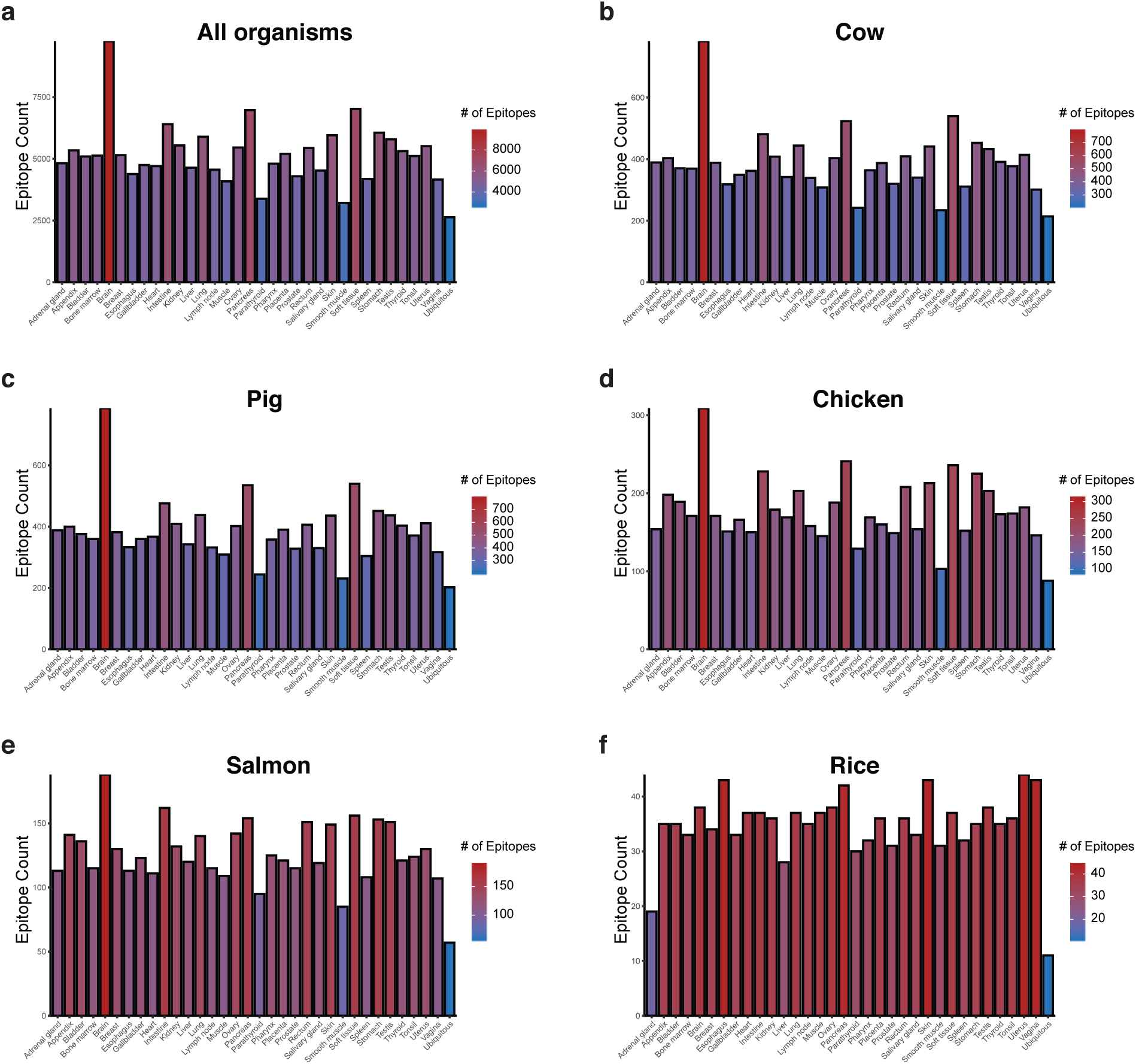
Tissue expression pattern of food-derived human autoimmune epitopes. Expression by organ of epitopes present in (**a**) all organisms commonly consumed as food analyzed, (**b**) cow, (**c**) pig, (**d**) chicken, (**e**) salmon, (**f**) rice.

Next, we sought to investigate the binding of the identified diet-derived autoimmune epitopes to HLA alleles. As per the International Immunogenetics Database, there are currently almost 20,000 different HLA class I and 10,000 HLA class II alleles identified in the world population, resulting in every person having a unique HLA repertoire. Therefore, mapping diet-derived epitope HLA binding has the potential to personalize the assessment of the risk that each food poses in terms of autoimmunogenicity. In addition, many HLA alleles have been either associated or found to be protective against multiple autoimmune disorders, allowing us to gain insight into the relative affinity of a given disease-linked HLA allele to diet-derived epitopes implicated in that same disease. HLA class I restricted epitopes are typically 8-11 amino acids long^26^. Yet, peptides as long as 15 amino acids have been found to bind to some HLA class I alleles with affinity and stability comparable to those of canonical length^27,28^. Moreover, some 7 amino acid long peptides can stably bind to HLA class I and, curiously, peptides as short as 4 amino acids can also occupy the HLA class I binding groove in pairs, creating neoepitopes and eliciting CD8^+^ T cell responses^29^. HLA class II restricted epitopes, on the other hand, usually span 13-20 amino acids in length^30,31^. Nevertheless, unlike HLA class I, there is no known limit to the number of amino acids an HLA class II epitope can have, with peptides up to 40 amino acids long reported as natural ligands of HLA class II alleles^32^.

We used an *in silico* MHC binding prediction tool^33^ (tools.immuneepitope.org) to interrogate 8-14 amino acid long and 11-30 amino acid long autoimmune epitopes binding to HLA class I and class II alleles, respectively. Of the 10,605 epitopes previously implicated in human autoimmune disorders, 2306 are potential HLA class I epitopes and 4547 are potential HLA class II epitopes, according to these thresholds. We focused our analysis on five common autoimmune diseases: T1D, RA, Behcet Syndrome, SLE, and MS. These diseases have a large number of implicated epitopes (“hot diseases”, > 500 epitopes) and several HLA alleles previously shown either to predispose individuals towards (odds ratio, OR > 1) or protect against (OR < 1) the development of these diseases (Figure 5).

**Figure 5.**
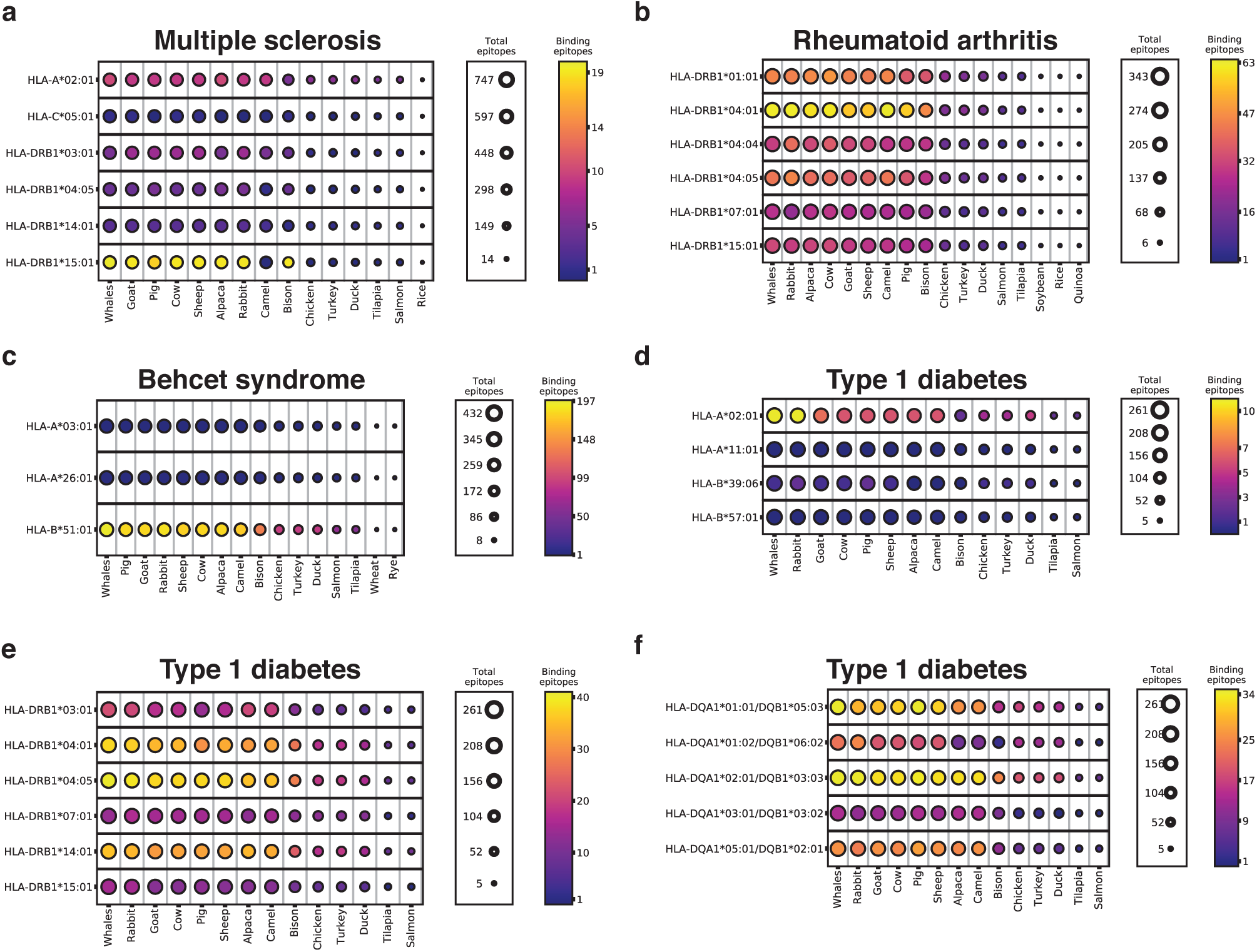
Differential binding of diet-derived autoimmune epitopes to disease-associated HLA alleles. Binding of epitopes implicated in four “hot” diseases, i.e. multiple sclerosis (**a**), rheumatoid arthritis (**b**), Behcet syndrome (**c**), and type 1 diabetes (**d**,**e**,**f**), to HLA alleles either associated with or protective against each disease was predicted *in silico* using tools.immuneepitope.org. Each circle represents the autoimmune epitopes present in a species commonly consumed as food associated with the disease (x axis). The size of the circle represents the number of total epitopes contained in the organism that have been associated with the disease (right hand “total epitopes” legend), whereas the color represents the number of those epitopes predicted to bind to each HLA allele along the y axis (right hand “binding epitopes” heatmap legend).

Overall, organisms with a higher autoimmune epitope content yielded more epitopes bound to the analyzed HLA alleles. Strikingly, HLA alleles associated with a given disease have more epitopes predicted to bind to them than those found to be protective against that same disease (Figure 5), consistent with the notion that some HLA alleles are protective because they don’t present pathogenic epitopes as frequently as other HLA alleles, especially those associated with the disease.

Indeed, for MS, HLA-DRB1*15:01 (associated^34,35^, OR = 3.10) is predicted to present up to 20 epitopes present in all red meat animals other than camel, whereas HLA-DRB1*14:01 (protective^35,36^, OR = 0.54) presents almost none (Figure 5a). For RA, HLA-DRB*04:01, an allele associated^37^ with the disease (OR = 3.3.1) presents the most peptides, whereas the protective^37^ alleles HLA-DRB*07:01 (OR = 0.66) and HLA-DRB*13:01 (OR = 0.58) present the least (Figure 5b). Of note, HLA-DRB*04:04 and HLA-DRB*04:05 are also risk alleles for RA^37^ (with OR of 2.80 and 3.27, respectively) but bind to visibly less diet-derived epitopes than HLA-DRB*04:01 (Figure 5b). Unlike most autoimmune diseases, Behcet syndrome has been mostly associated with HLA class I alleles^38,39^. In agreement with human genetic associations, we found that HLA-B*51:01(OR = 5.78) is predicted to bind a significant number of disease epitopes, whereas HLA-A*03:01 (OR = 0.6) is not (Figure 5c).

Finally, we investigated diet-derived epitope binding to three categories of HLA alleles relevant in T1D biology: HLA-A and HLA-B, HLA-DRB, and HLA-DQA/HLA-DQB. With regards to class I alleles, the T1D-associated allele^40^ HLA-A*02:01 (OR = 1.35) bound the most epitopes, with whale and rabbit containing dramatically more binding autoimmune epitopes than other red meat sources. Unexpectedly, HLA-B*39:06 and HLA-B*57:01 have vastly different OR^40^ (10.30 and 0.19, respectively), yet neither bind diet-derived T1D epitopes (Figure 5d). As for class II alleles, epitope binding also does not always follow a pattern consistent with the concept that associated HLA alleles bind disease epitopes, whereas protective alleles do not: while neither HLA-DRB1*15:01 nor HLA-DQA1*01:02/-DQB1*06:02 bind to diet-derived T1D epitopes, agreeing with the OR of 0.03 for the HLA-DRB1*15:01–HLA-DQA1*01:02–HLA-DQB1*06:02 haplotype^40^, HLA-DRB1*14:01 and HLA-DQA1*01:01/-DQB1*05:03 bind to many epitopes, yet the OR for the corresponding haplotype^40^ is 0.02 (Figure 5e,f). Nevertheless, for three haplotypes associated with the disease^40^, HLA-DRB1*04:05–HLA-DQA1*03:01–HLA-DQB1*03:02 (OR = 11.37), HLA-DRB1*04:01–HLA-DQA1*03:01–HLA-DQB1*03:02 (OR = 8.39), and HLA-DRB1*03:01–HLA-DQA1*05:01–HLA-DQB1*02:01 (OR = 3.64), at least one of the HLA alleles was found to bind a large number of T1D epitopes (Figure 5e,f).

These observations led us to develop a personalized scorecard to determine food autoimmunogenicity for individual patients – the Gershteyn-Ferreira sensitivity passport. As a proof-of-principle, we analyzed 6 (out of a theoretical maximum of 9) pairs of HLA alleles taken from a pool of de-identified individual’s haplotypes for binding to autoimmune epitopes found in different organisms commonly consumed as food, and determined which diseases most of those epitopes have been implicated in. Focusing on the “hot” diseases (> 500 epitopes), we found that HLA-B*38:06 and HLA-C*12:02 only bind robustly to Behcet syndrome epitopes. HLA-A*02:01 binds to MS and T1D epitopes, as expected, yet mostly only to those found in whale and rabbit in the case of T1D. For HLA class II, all analyzed alleles bind to MS epitopes, with HLA-DRB1*04:01 also binding to RA and T1D epitopes (Figure 6). Inspecting binding to epitopes found in “warm” diseases (between 50 and 500 epitopes), we found that HLA-DRB1*04:01 binds to epitopes from myasthenia gravis, Sjogren’s syndrome and systemic scleroderma, whereas HLA-DRB1*03:01 and HLA-DRB1*15:01 are only likely to present systemic scleroderma epitopes (Extended Data Figure 4).

**Figure 6.**
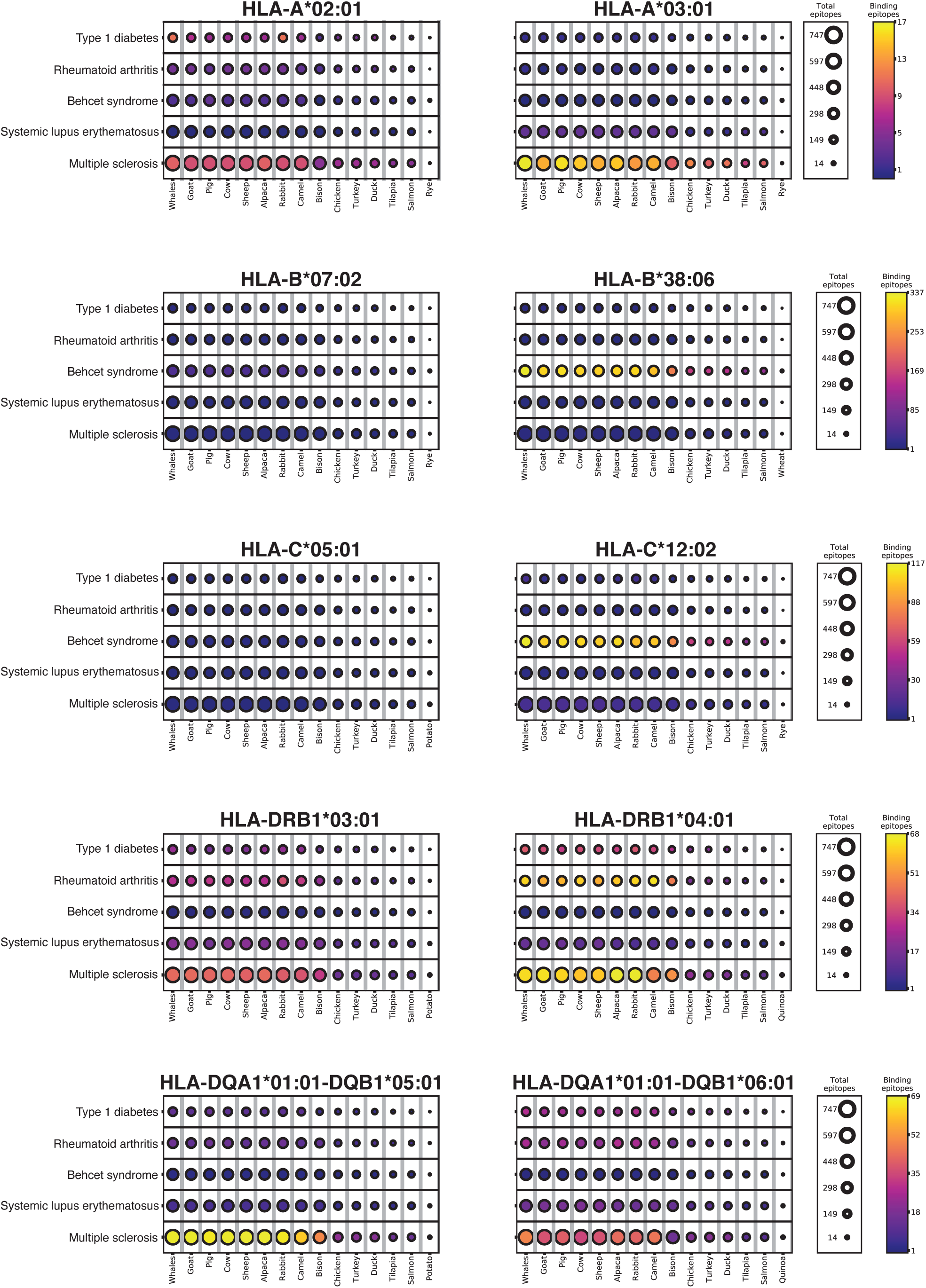
Gershteyn-Ferreira sensitivity passport. Binding of total autoimmune epitopes present in food to ten HLA alleles (six HLA class I and four HLA class II alleles, with HLA-DQA and HALA-DQB being combined – alpha and beta chain of the HLA-DQ molecule) from one individual was predicted *in silico* using tools.immuneepitope.org. Each circle represents the autoimmune epitopes present in a species commonly consumed as food (x axis) implicated in a human autoimmune disease (y axis). The size of the circle represents the number of total epitopes contained in the organism that have been associated with the disease (right hand “total epitopes” legend), whereas the color represents the number of those epitopes predicted to bind to the HLA allele (right hand “binding epitopes” heatmap legend).

## Discussion

Autoimmunity has been on the rise around the globe at a fast pace, leading many to believe that environmental factors, not genetics, are mostly responsible for this trend. Processed foods play a large role in the American diet and have been shown to compromise intestinal barrier integrity^17^. In the United States, while processed meat consumption has remained stable over the past two decades, total meat consumption has decreased^41^. Therefore, the proportion of processed meat in American diets has increased.

Celiac disease, an autoimmune disease where the lining of the small intestine is destroyed, has a well-defined interaction with diet and specific HLA alleles. Partly undigested gluten peptides cross the intestinal barrier and are modified by tissue transglutaminase 2 (TG2), which generates epitopes that are then presented to autoreactive gluten-specific T cells^20,42^. Over 80% of celiac disease patients carry HLA-DQ2 (HLA-DQA1*05:01/-DQB1*02:01, OR = 15.1)^43,44^. Gluten-specific regulatory T cells (Tregs), a subset of T lymphocytes dedicated to maintaining homeostasis by limiting immune responses^45^, have been found to have a defective suppressive capacity in celiac disease patients^46^. Removing gluten from the diet leads to remission of the disease. Can we work towards determining the impact of diet on other autoimmune diseases with the same degree of mechanistic detail? We predict that foods, containing epitopes implicated in a certain autoimmune disease, can be triggers to exacerbate its symptoms upon frequent consumption.

One limitation of the present study lies in the fact that only unmodified linear peptide epitopes were considered. Hence, post-translational modifications, such as citrullination^47^, and hybrid peptides, such as the hybrid insulin peptides (HIP) created by linkage of an insulin fragment to another peptide in beta cells^48^, previously shown to be implicated in autoimmunity, were not analyzed. Moreover, food production and consumption practices can significantly affect the specific protein content of the final product. For instance, proteins found only in the breast muscle of fast-growth chickens have been identified^49^, whereas cooking food can alter their antigen composition and their impact on the gut microbiome^50^. In turn, an individual’s microbiome composition can determine his or her propensity to intestinal inflammation and autoimmunity^51^, adding an additional layer warranting future investigations. Finally, we predict that epidemiological studies will reveal differences in the incidence and/or severity of certain autoimmune disorders in populations with specific dietary habits, namely exclusion of one or more foods, and that studies using humanized animals expressing different HLA alleles subjected to dietary antigen exposure regimens will refine our mechanistic understanding of the interaction between diet and autoimmunity.

In summary, we systematically dissected the autoimmunogenicity of 24 organisms, 22 of which are commonly consumed as food. We mapped not only their content of epitopes previously implicated in human autoimmune disorders (Figure 1), but also assessed the capacity of different HLA alleles to present these peptides. This research revealed striking differences in binding, hence potential food autoimmunogenicity, across HLA alleles (Figure 5), an important step towards personalizing this information for each autoimmune disease patient (Gershteyn-Ferreira Sensitivity Passport, Figure 6). Overall, our platform will help shed light on the often observed yet seldom understood impact of diet on autoimmune disorders.

## Methods

### Human autoimmune epitope identification in food sources

To identify epitopes implicated in human autoimmune disease prevalent in commonly consumed foods, we aggregated 10,605 linear peptide epitopes implicated in 70 autoimmune diseases, available^23^ at www.iedb.org (retrieved September 10, 2019), and cross-referenced them with the proteomes of 24 organisms: alpaca (taxid:30538), bat (taxid:9397), bison (taxid:9901), salmon (taxid:8030), camel (taxid:419612), whale (taxid:9721), chicken (taxid:9031), chimpanzee (taxid:9598), cow (taxid:9913), duck (taxid:8839), human (Homo sapiens, taxid:9606), goat (taxid:9925), rice (taxid:39947), lettuce (taxid:4236), turkey (taxid:9102), tilapia (taxid:8128), rabbit (taxid:9986), pig (taxid:9823), potato (taxid:4113), quinoa (taxid:63459), rye (taxid:4550), sheep (taxid:9940), soybean (taxid:3847), and wheat (taxid:4564). We then utilized Node.js, AWS EC2, and PostgreSQL database technologies to automate data gathering and expedite analysis. Data gathering automation was broken into two custom processes. Process #1 queried the National Center for Biotechnology Information (NCBI) blastp system. The query request consisted of the unique epitope ID and the list of organisms. All the queries were checked against the NCBI refseq_protein database. The API supports a list of organisms as a valid query criterion, significantly cutting down the round trip of each query. Once a request was made for a particular epitope in the database, Process #1 created a record with the Request ID where the uniqueness of the record was the epitope ID and the organism. The record was marked as pending to indicate that it is ready to be retrieved for processing by Process #2. The process continued until all the epitopes had been queried and marked as pending. Process #2 retrieved the results of the queries via Request ID from Process #1. As the query could take an uncertain amount of time to finish, the process iterated through the list of all the records that had a valid Request ID to obtain the results. In the query result data, the top hit for each organism was selected and then, if the Query Coverage and Identity Percentage values both equaled 100, the record result was considered as a “hit” (total match). In all other cases, a “miss” was recorded. Once hit or miss was recorded, the process was marked as complete so the automated query would ignore on the next pass of requests. The system continued polling NCBI servers until all results were gathered. Once both processes were fully executed, we had compiled a fully comprehensive database of autoimmune human-organism epitope overlap. An interactive website for researchers and the public to explore this database will be available at www.immunodietica.com.

### Mapping human autoimmune epitope tissue expression

A list of 10,605 epitopes associated with human autoimmune diseases were passed into a function converting them into FASTA format and queried against the Swiss-Prot database (hosted by NIH) to acquire the UniProt name associated with the protein containing each epitope in the list. Queries were performed by directly parsing through the database. In the instance where a direct lookup failed, BLASTp (ncbi-blast-2.10.0+) was run to find a match within the same database. Due to the lengthy runtime and computational intensity associated with BLASTp queries, BLASTp was only run if a linear scan of the database yielded zero results. Threshold parameters were used to limit queries to only one result and only perfect matches (100% identity across all amino acids in query and matching sequence). The gene symbol for each epitope was obtained by querying db2db API (bioDBnet) with the associated UniProt obtained in the previous query. This query resulted in a data table linking the original list of epitopes with their associated gene symbols. Expression profiles for proteins in human tissues were obtained from the Human Protein Atlas (data from the Human Protein Atlas version 19.3 and Ensembl version 92.38). For each protein in the tissue expression data, Ensembl name, gene symbol name, and tissue and cellular localization were provided. In addition, the data detailed the level of expression (High, Medium, Low, Not detected) and the reliability of the described expression level and localization based on existing literature (Approved, Enhanced, Supported, Uncertain). To obtain the tissue-level localization of the autoimmune disease-associated epitopes, the tissue expression data from the Human Protein Atlas and the list of epitopes were joined together using gene symbol as a matching identity key. Data for proteins in which reliability was “Uncertain” or expression level was “Undetected” were removed. A separate table labeling each tissue with its associated organ and tissue system was joined in order to provide tissue-system and organ-level localization of the epitopes. Each epitope in the resulting data table was labeled with its associated autoimmune disease. This resulted in a database, plotted and shown in Figure 3. For Figure 4, the same 10,605 epitopes were queried against the complete proteome of the 24 organisms indicated above in *Human autoimmune epitope identification in food sources*. Only matches with 100% identity were retained. The same pipeline detailed above was followed and the final table of proteins and their organ-level expression was matched against the complete set of diet-derived epitopes generated in the previous step. The resulting database consisted of the amino acid sequence, gene symbol, associated organism, and organ-level expression and localization for each epitope. The database was then filtered for cow (taxid:9913), pig (taxid:9823), chicken (taxid:9031), salmon (taxid:8030), and rice (taxid:39947).

### HLA binding prediction

The IEDB MHC-I and MHC-II Binding Prediction tools (tools.immuneepitope.org) API were used to estimate binding affinity of each autoimmune epitope to various HLA alleles *in silico*^33^. For HLA class I alleles, 2306 epitopes were assessed, whereas for HLA class II, 4547 epitopes were analyzed. For epitopes of length greater than 15 amino acids, sliding 15-mer peptides were queried. The length of each queried peptide was set to 8-14 amino acids for each HLA class I allele and 11-30 amino acids for each HLA class II allele. As per previous protocols^33^, epitope-HLA pairs with percentile rank scores greater than 1% were considered strong binders to HLA class I and those with percentile rank scores greater than 10% were considered strong binders to HLA class II, respectively, and used for downstream analysis. BLASTp was used to identify organisms commonly used as food sources that also had these strong-binding epitopes. The number of binding epitopes for each associated organism and type of autoimmune disease were then recorded to create a map of predicted food sensitivity (Gershteyn-Ferreira sensitivity passport) for the set of HLA alleles originally queried.

## Acknowledgements

This work was funded by Ajax Biomedical Foundation (Newton, MA). L.M.R.F. is the Jeffrey G. Klein Diabetes Fellow. We thank our colleagues for their support: Dr. Victor Goldmacher, Dr. Andrey Vyshedskiy, Mikhail Gershteyn, Arkady Gershteyn.

---

**Extended Data Figure 1.**
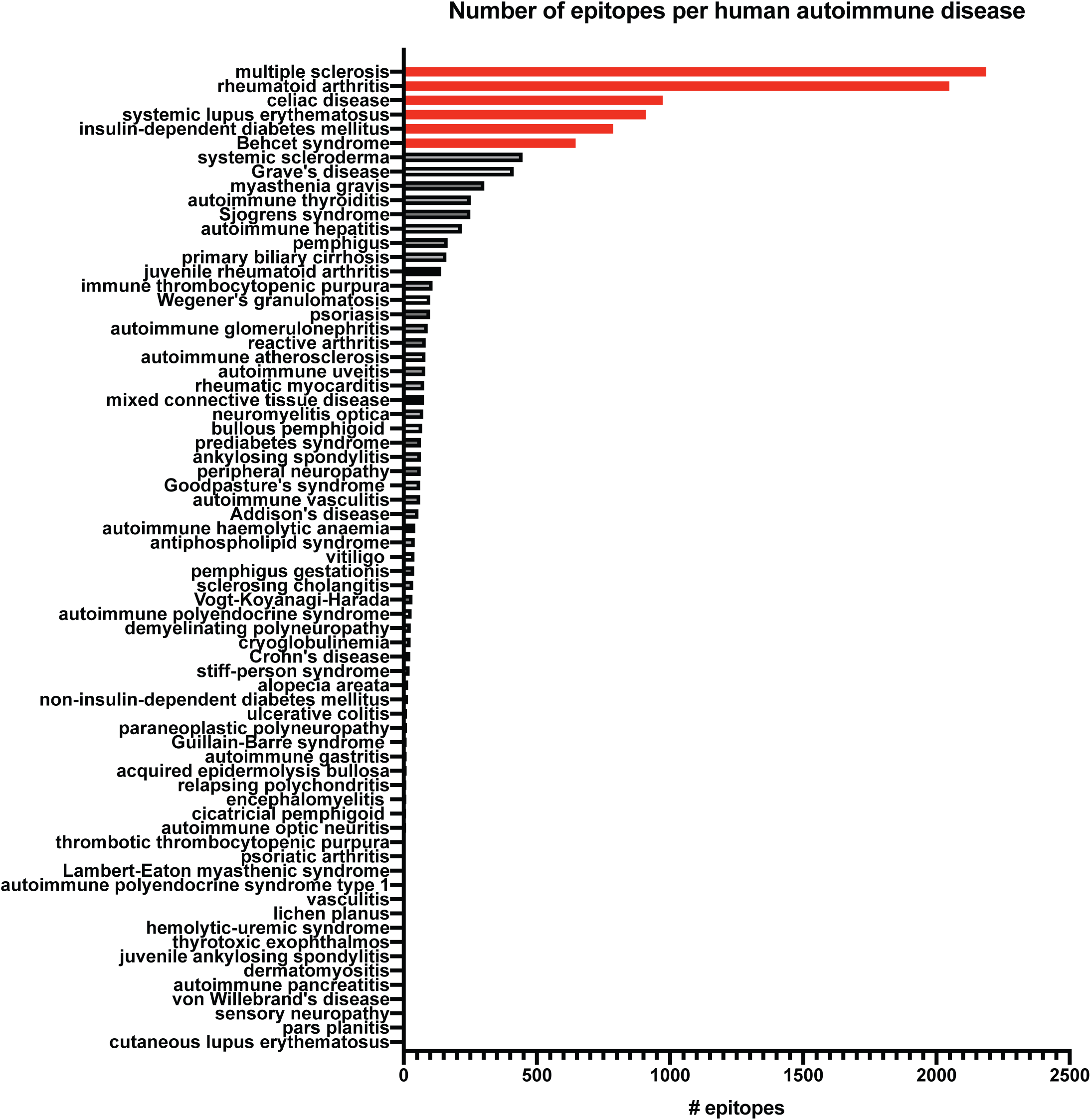
Number of epitopes implicated in human autoimmune diseases. Number of linear peptide epitopes identified per disease. Diseases ordered in decreasing number of epitopes known from top to bottom. The six with most epitopes identified are colored in red.

**Extended Data Figure 2.**
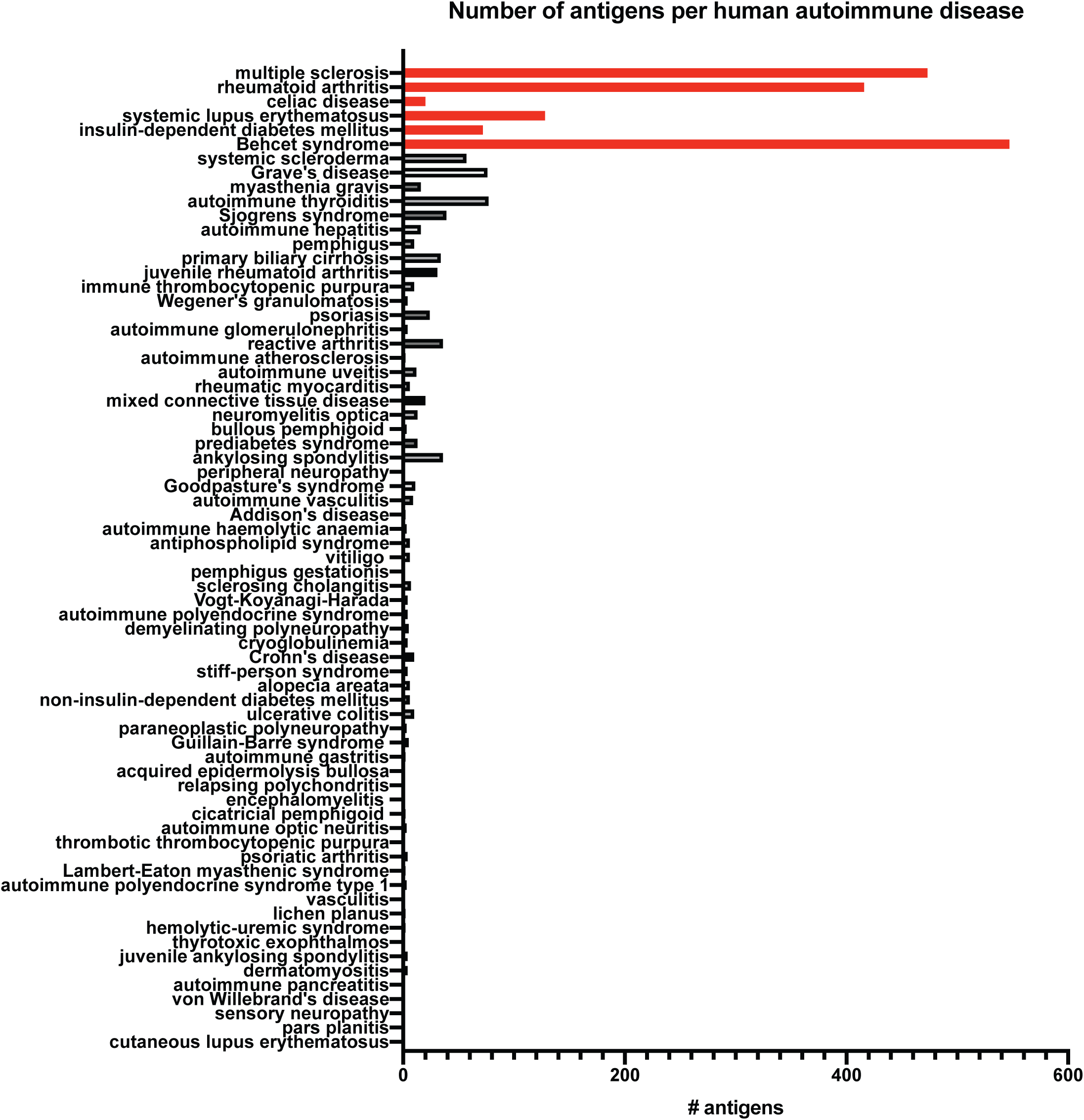
Number of antigens implicated in human autoimmune diseases. Number of antigens identified per disease. Diseases ordered in decreasing number of epitopes known from top to bottom. The six with most epitopes identified are colored in red.

**Extended Data Fig 3.**
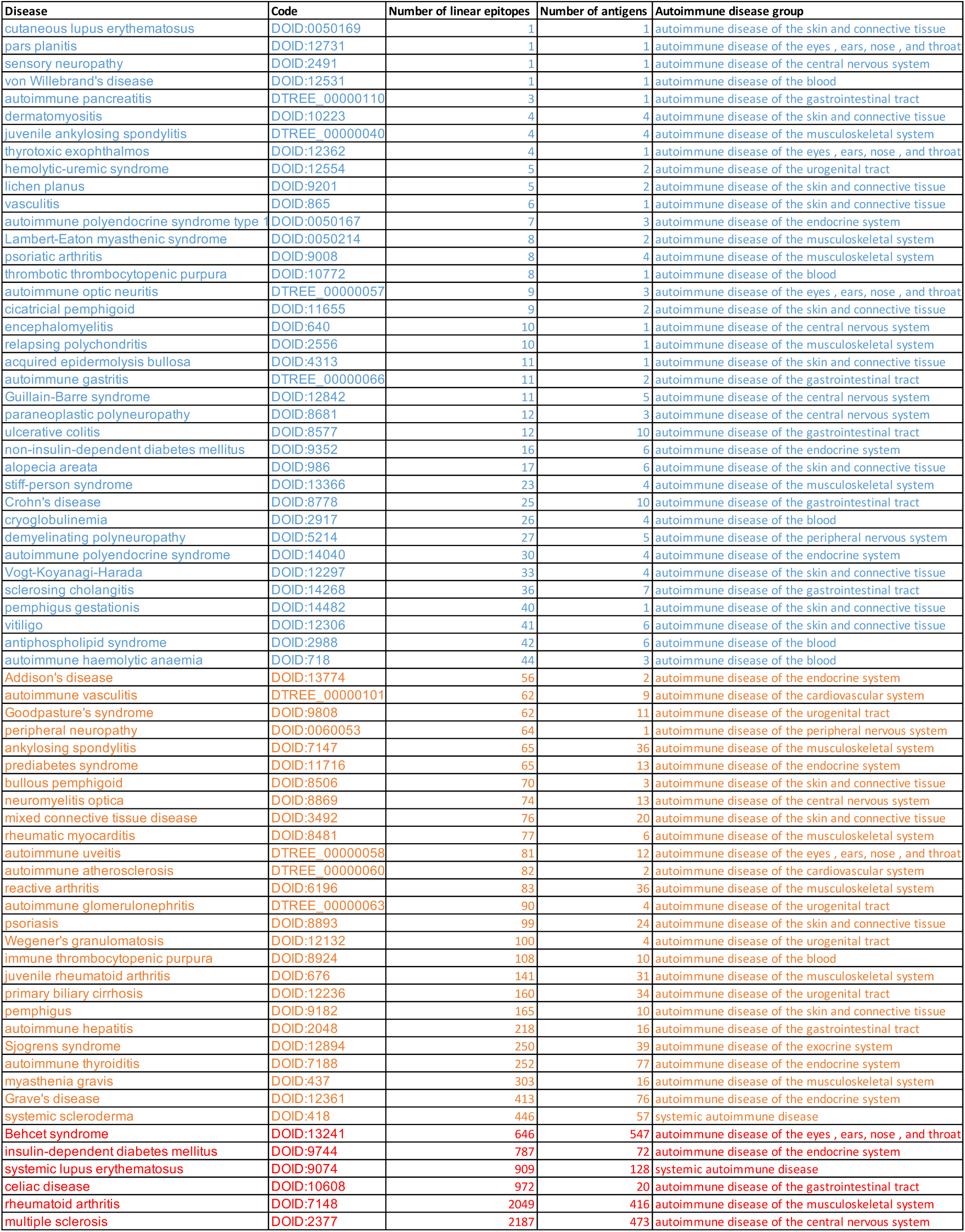
Classification of autoimmune diseases based on number of identified epitopes. Human autoimmune disorders ordered and grouped according to the number of known implicated linear peptide epitopes, retrieved from www.iedb.org (September 10, 2019). Color code: red, “hot” (> 500); orange, “warm” (50-500); blue, “cold” (< 50).

**Extended Data Fig 4.**
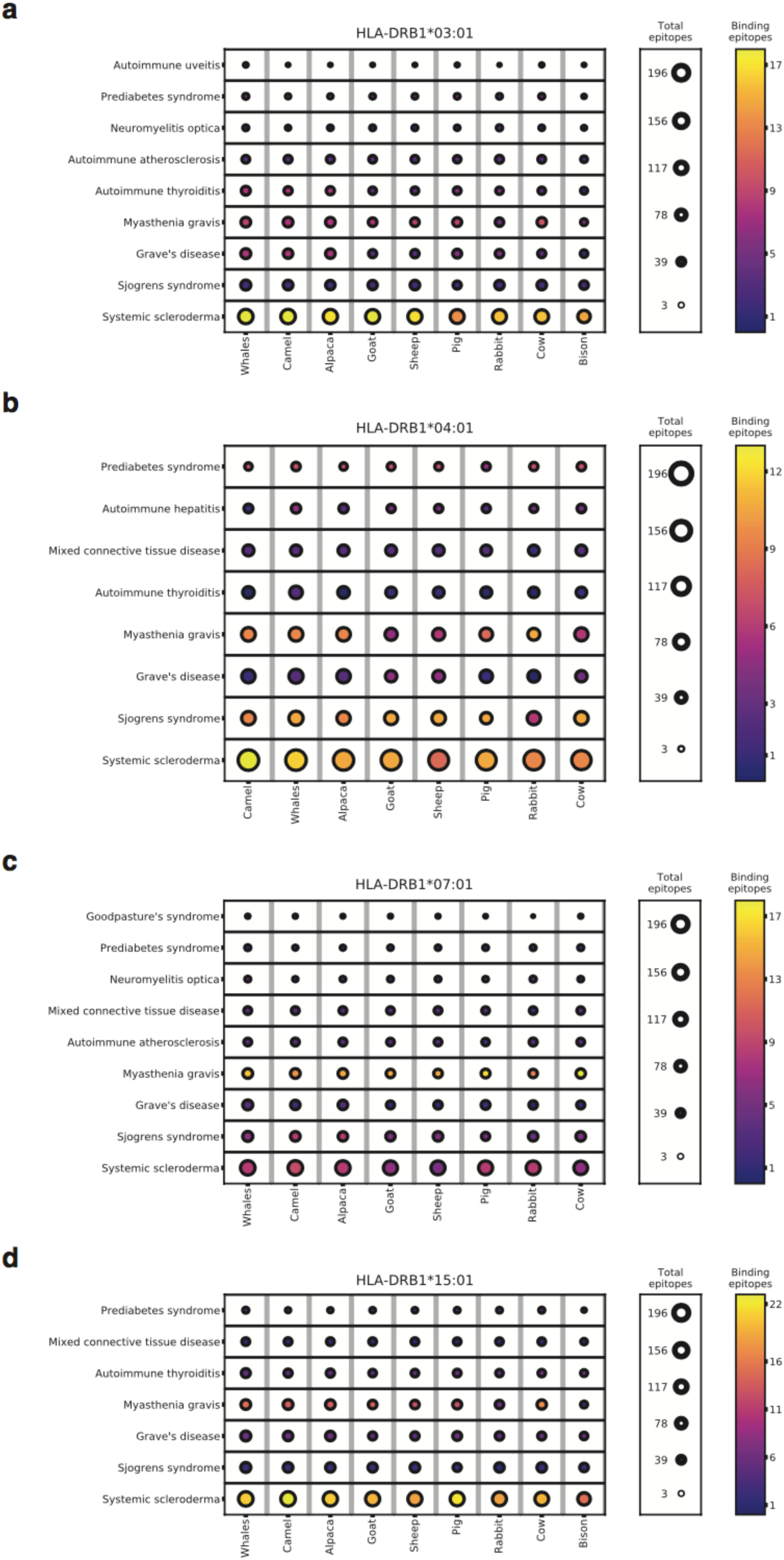
Differential binding of disease-specific epitopes to HLA alleles. Predicted binding (www.tools.immunepitope.org) of epitopes implicated in “warm” autoimmune diseases (50-500 epitopes) to HLA-DRB1*03:01 **(a)**, HLA-DRB1*04:01 **(b)**, HLA-DRB1*07:01 **(c)**, and HLA-DRB1*15:01 **(d)**.

## References

1 Barabási, A., Menichetti, G. & Loscalzo, J. The unmapped chemical complexity of our diet. Nat Food 1, 33–37, doi: https://doi.org/10.1038/s43016-019-0005-1 (2020).

2 Cooper, G. S., Miller, F. W. & Pandey, J. P. The role of genetic factors in autoimmune disease: implications for environmental research. Environ Health Perspect 107 Suppl 5, 693–700, doi: 10.1289/ehp.99107s5693 (1999).

3 Vojdani, A. A Potential Link between Environmental Triggers and Autoimmunity. Autoimmune Dis 2014, 437231, doi: 10.1155/2014/437231 (2014).

4 Malosse, D. & Perron, H. Correlation analysis between bovine populations, other farm animals, house pets, and multiple sclerosis prevalence. Neuroepidemiology 12, 15–27, doi: 10.1159/000110295 (1993).

5 Kjeldsen-Kragh, J. et al. Controlled trial of fasting and one-year vegetarian diet in rheumatoid arthritis. Lancet 338, 899–902 (1991).

6 Bock, S. A. & Atkins, F. M. Patterns of food hypersensitivity during sixteen years of double-blind, placebo-controlled food challenges. J Pediatr 117, 561–567, doi: 10.1016/s0022-3476(05)80689-4 (1990).

7 Kleinewietfeld, M. et al. Sodium chloride drives autoimmune disease by the induction of pathogenic TH17 cells. Nature 496, 518–522, doi: 10.1038/nature11868 (2013).

8 Wu, C. et al. Induction of pathogenic TH17 cells by inducible salt-sensing kinase SGK1. Nature 496, 513–517, doi: 10.1038/nature11984 (2013).

9 Kawanishi, K. et al. Human species-specific loss of CMP-N-acetylneuraminic acid hydroxylase enhances atherosclerosis via intrinsic and extrinsic mechanisms. Proc Natl Acad Sci U S A 116, 16036–16045, doi: 10.1073/pnas.1902902116 (2019).

10 Rojas, M. et al. Molecular mimicry and autoimmunity. J Autoimmun 95, 100–123, doi: 10.1016/j.jaut.2018.10.012 (2018).

11 Gough, S. C. & Simmonds, M. J. The HLA Region and Autoimmune Disease: Associations and Mechanisms of Action. Curr Genomics 8, 453–465, doi: 10.2174/138920207783591690 (2007).

12 Shahrizaila, N. & Yuki, N. Guillain-barre syndrome animal model: the first proof of molecular mimicry in human autoimmune disorder. J Biomed Biotechnol 2011, 829129, doi: 10.1155/2011/829129 (2011).

13 Gil-Cruz, C. et al. Microbiota-derived peptide mimics drive lethal inflammatory cardiomyopathy. Science 366, 881–886, doi: 10.1126/science.aav3487 (2019).

14 Hvatum, M., Kanerud, L., Hallgren, R. & Brandtzaeg, P. The gut-joint axis: cross reactive food antibodies in rheumatoid arthritis. Gut 55, 1240–1247, doi: 10.1136/gut.2005.076901 (2006).

15 Lachance, D. H. et al. An outbreak of neurological autoimmunity with polyradiculoneuropathy in workers exposed to aerosolised porcine neural tissue: a descriptive study. Lancet Neurol 9, 55–66, doi: 10.1016/S1474-4422(09)70296-0 (2010).

16 Bischoff, S. C. et al. Intestinal permeability--a new target for disease prevention and therapy. BMC Gastroenterol 14, 189, doi: 10.1186/s12876-014-0189-7 (2014).

17 Lerner, A. & Matthias, T. Changes in intestinal tight junction permeability associated with industrial food additives explain the rising incidence of autoimmune disease. Autoimmun Rev 14, 479–489, doi: 10.1016/j.autrev.2015.01.009 (2015).

18 Purohit, V. et al. Alcohol, intestinal bacterial growth, intestinal permeability to endotoxin, and medical consequences: summary of a symposium. Alcohol 42, 349–361, doi: 10.1016/j.alcohol.2008.03.131 (2008).

19 Mu, Q., Kirby, J., Reilly, C. M. & Luo, X. M. Leaky Gut As a Danger Signal for Autoimmune Diseases. Front Immunol 8, 598, doi: 10.3389/fimmu.2017.00598 (2017).

20 Verdu, E. F. & Danska, J. S. Common ground: shared risk factors for type 1 diabetes and celiac disease. Nat Immunol 19, 685–695, doi: 10.1038/s41590-018-0130-2 (2018).

21 Erickson, M. A. & Banks, W. A. Age-Associated Changes in the Immune System and Blood(-)Brain Barrier Functions. Int J Mol Sci 20, doi: 10.3390/ijms20071632 (2019).

22 Jarius, S. et al. Mechanisms of disease: aquaporin-4 antibodies in neuromyelitis optica. Nat Clin Pract Neurol 4, 202–214, doi: 10.1038/ncpneuro0764 (2008).

23 Vita, R. et al. The Immune Epitope Database (IEDB): 2018 update. Nucleic Acids Res 47, D339–D343, doi: 10.1093/nar/gky1006 (2019).

24 Rowntree, L. C. et al. Inability To Detect Cross-Reactive Memory T Cells Challenges the Frequency of Heterologous Immunity among Common Viruses. J Immunol 200, 3993–4003, doi: 10.4049/jimmunol.1800010 (2018).

25 Vojdani, A. Reaction of food-specific antibodies with different tissue antigens. Int J Food Sci Technol 55, 1800–1815 (2019).

26 Trolle, T. et al. The Length Distribution of Class I-Restricted T Cell Epitopes Is Determined by Both Peptide Supply and MHC Allele-Specific Binding Preference. J Immunol 196, 1480–1487, doi: 10.4049/jimmunol.1501721 (2016).

27 Hassan, C. et al. Naturally processed non-canonical HLA-A*02:01 presented peptides. J Biol Chem 290, 2593–2603, doi: 10.1074/jbc.M114.607028 (2015).

28 Remesh, S. G. et al. Unconventional Peptide Presentation by Major Histocompatibility Complex (MHC) Class I Allele HLA-A*02:01: BREAKING CONFINEMENT. J Biol Chem 292, 5262–5270, doi: 10.1074/jbc.M117.776542 (2017).

29 Xiao, Z., Ye, Z., Tadwal, V. S., Shen, M. & Ren, E. C. Dual non-contiguous peptide occupancy of HLA class I evoke antiviral human CD8 T cell response and form neo-epitopes with self-antigens. Sci Rep 7, 5072, doi: 10.1038/s41598-017-05171-w (2017).

30 Stern, L. J. et al. Crystal structure of the human class II MHC protein HLA-DR1 complexed with an influenza virus peptide. Nature 368, 215–221, doi: 10.1038/368215a0 (1994).

31 O’Brien, C., Flower, D. R. & Feighery, C. Peptide length significantly influences in vitro affinity for MHC class II molecules. Immunome Res 4, 6, doi: 10.1186/1745-7580-4-6 (2008).

32 van Lummel, M. et al. Dendritic Cells Guide Islet Autoimmunity through a Restricted and Uniquely Processed Peptidome Presented by High-Risk HLA-DR. J Immunol 196, 3253–3263, doi: 10.4049/jimmunol.1501282 (2016).

33 Fleri, W. et al. The Immune Epitope Database and Analysis Resource in Epitope Discovery and Synthetic Vaccine Design. Front Immunol 8, 278, doi: 10.3389/fimmu.2017.00278 (2017).

34 Patsopoulos, N. A. et al. Fine-mapping the genetic association of the major histocompatibility complex in multiple sclerosis: HLA and non-HLA EFects. PLoS Genet 9, e1003926, doi: 10.1371/journal.pgen.1003926 (2013).

35 Hollenbach, J. A. & Oksenberg, J. R. The immunogenetics of multiple sclerosis: A comprehensive review. J Autoimmun 64, 13–25, doi: 10.1016/j.jaut.2015.06.010 (2015).

36 Toro, J. et al. HLA-DRB1*14 is a protective allele for multiple sclerosis in an admixed Colombian population. Neurol Neuroimmunol Neuroinflamm 3, e192, doi: 10.1212/NXI.0000000000000192 (2016).

37 Balandraud, N. et al. HLA-DRB1 genotypes and the risk of developing anti citrullinated protein antibody (ACPA) positive rheumatoid arthritis. PLoS One 8, e64108, doi: 10.1371/journal.pone.0064108 (2013).

38 Hughes, T. et al. Identification of multiple independent susceptibility loci in the HLA region in Behcet’s disease. Nat Genet 45, 319–324, doi: 10.1038/ng.2551 (2013).

39 Ombrello, M. J. et al. Behcet disease-associated MHC class I residues implicate antigen binding and regulation of cell-mediated cytotoxicity. Proc Natl Acad Sci U S A 111, 8867–8872, doi: 10.1073/pnas.1406575111 (2014).

40 Noble, J. A. & Valdes, A. M. Genetics of the HLA region in the prediction of type 1 diabetes. Curr Diab Rep 11, 533–542, doi: 10.1007/s11892-011-0223-x (2011).

41 Zeng, L. et al. Trends in Processed Meat, Unprocessed Red Meat, Poultry, and Fish Consumption in the United States, 1999-2016. J Acad Nutr Diet 119, 1085–1098 e1012, doi: 10.1016/j.jand.2019.04.004 (2019).

42 Molberg, O. et al. Tissue transglutaminase selectively modifies gliadin peptides that are recognized by gut-derived T cells in celiac disease. Nat Med 4, 713–717 (1998).

43 Mubarak, A. et al. Human leukocyte antigen DQ2.2 and celiac disease. J Pediatr Gastroenterol Nutr 56, 428–430, doi: 10.1097/MPG.0b013e31827913f9 (2013).

44 Khosravi, A. et al. The likelihood ratio and frequency of DQ2/DQ8 haplotypes in Iranian patients with celiac disease. Gastroenterol Hepatol Bed Bench 9, 18–24 (2016).

45 Ferreira, L. M. R., Muller, Y. D., Bluestone, J. A. & Tang, Q. Next-generation regulatory T cell therapy. Nat Rev Drug Discov 18, 749–769, doi: 10.1038/s41573-019-0041-4 (2019).

46 Cook, L. et al. Circulating gluten-specific FOXP3(+)CD39(+) regulatory T cells have impaired suppressive function in patients with celiac disease. J Allergy Clin Immunol 140, 1592–1603 e1598, doi: 10.1016/j.jaci.2017.02.015 (2017).

47 Mathsson, L. et al. Antibodies against citrullinated vimentin in rheumatoid arthritis: higher sensitivity and extended prognostic value concerning future radiographic progression as compared with antibodies against cyclic citrullinated peptides. Arthritis Rheum 58, 36–45, doi: 10.1002/art.23188 (2008).

48 Wiles, T. A. et al. Identification of Hybrid Insulin Peptides (HIPs) in Mouse and Human Islets by Mass Spectrometry. J Proteome Res 18, 814–825, doi: 10.1021/acs.jproteome.8b00875 (2019).

49 Phongpa-Ngan, P., Grider, A., Mulligan, J. H., Aggrey, S. E. & Wicker, L. Proteomic analysis and differential expression in protein extracted from chicken with a varying growth rate and water-holding capacity. J Agric Food Chem 59, 13181–13187, doi: 10.1021/jf202622n (2011).

50 Carmody, R. N. et al. Cooking shapes the structure and function of the gut microbiome. Nat Microbiol 4, 2052–2063, doi: 10.1038/s41564-019-0569-4 (2019).

51 Zitvogel, L. & Kroemer, G. Immunostimulatory gut bacteria. Science 366, 1077–1078, doi: 10.1126/science.aaz7595 (2019).

