## Supplementary figures and images for "Immunodietica: interrogating the role of diet in autoimmune disease"

### Extended Data Figure 1

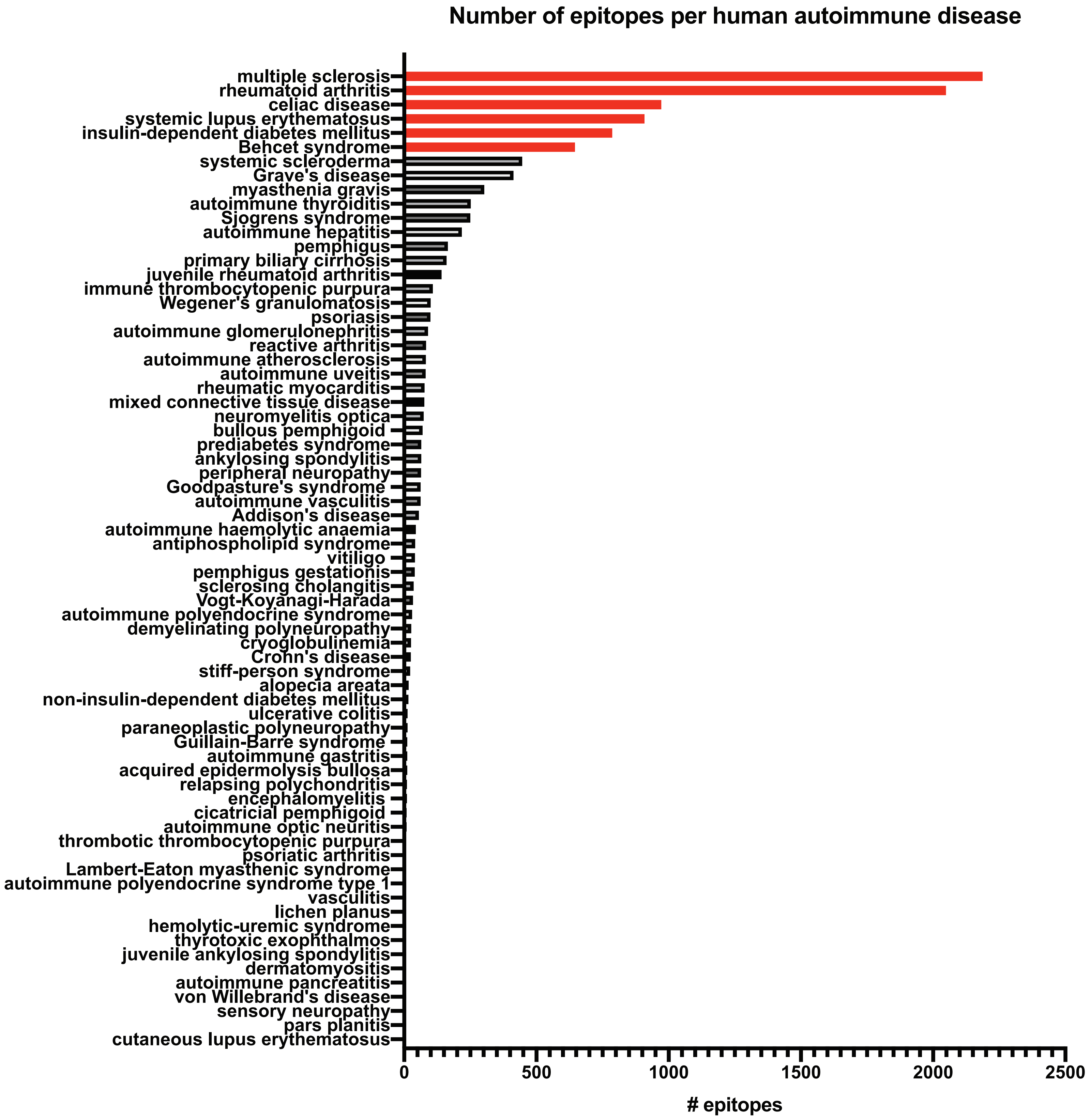

### Extended Data Figure 2

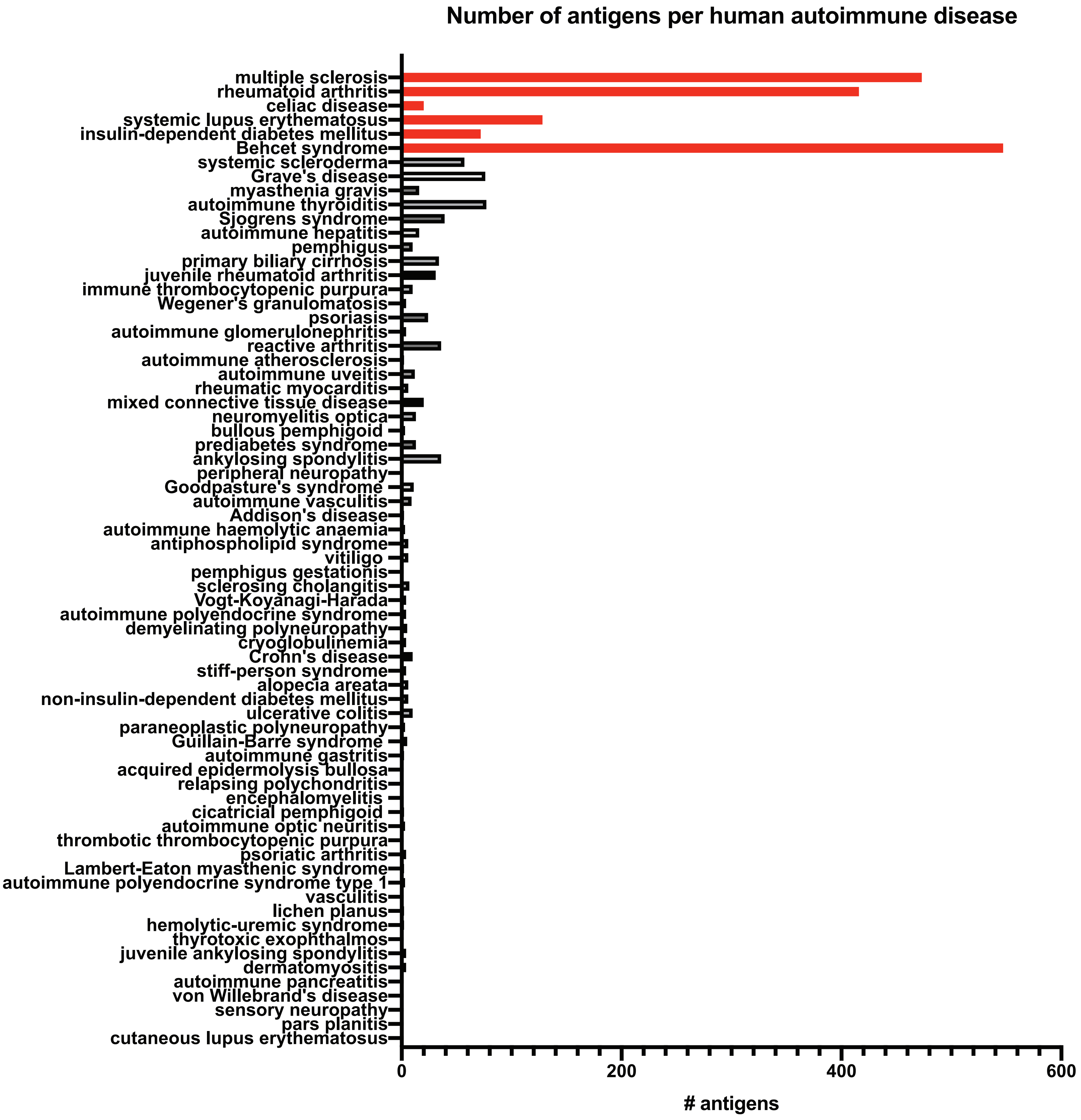

### Extended Data Figure 4

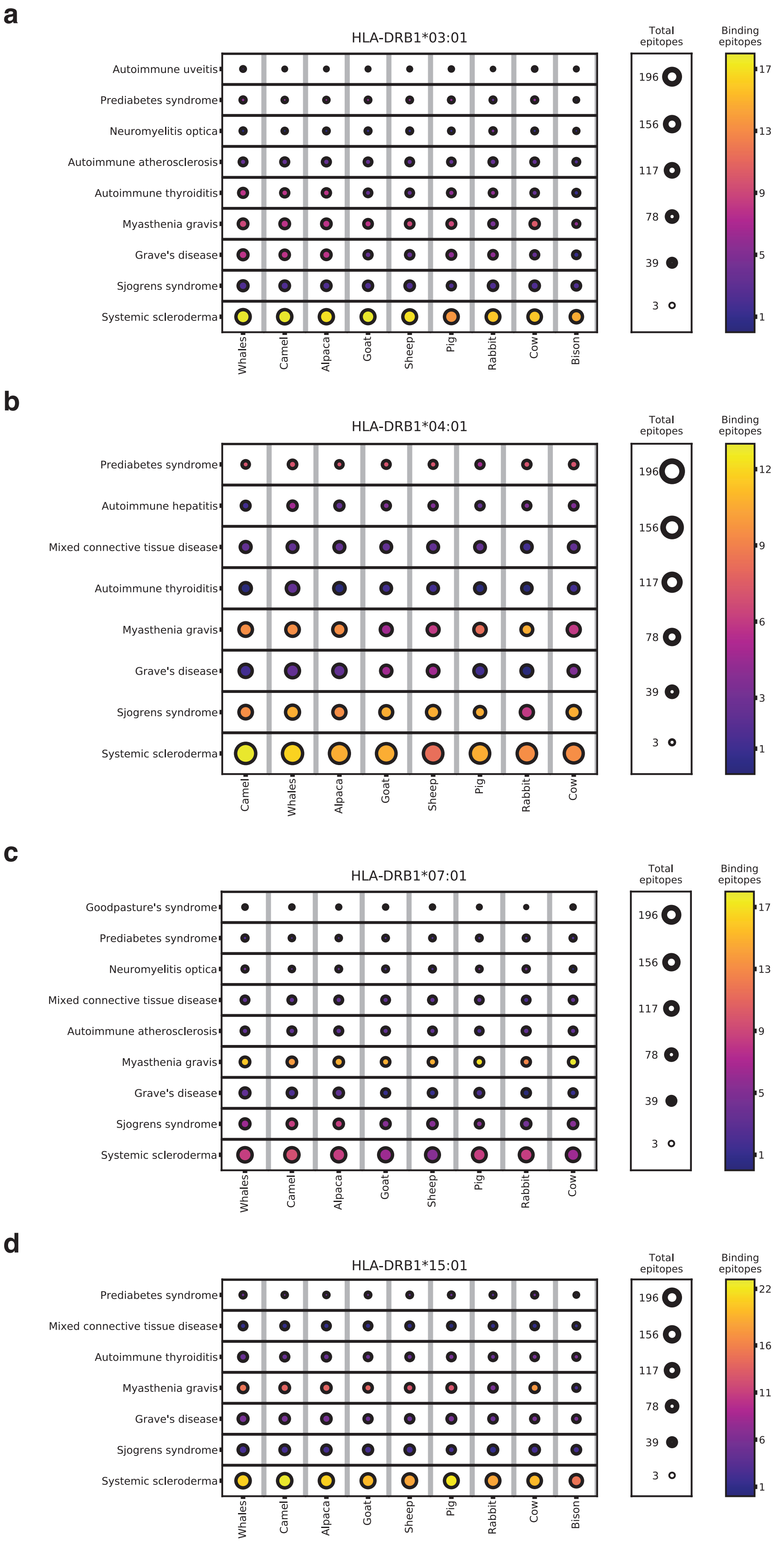
