## Extended Data Figure 3 for "Immunodietica: interrogating the role of diet in autoimmune disease"

| Disease | Code | Number of linear epitopes | Number of antigens | Autoimmune disease group |
|---|---|---|---|---|
| cutaneous lupus erythematosus | DOID:0050169 | 1 | 1 | autoimmune disease of the skin and connective tissue |
| pars planitis | DOID:12731 | 1 | 1 | autoimmune disease of the eyes , ears, nose , and throat |
| sensory neuropathy | DOID:2491 | 1 | 1 | autoimmune disease of the central nervous system |
| von Willebrand's disease | DOID:12531 | 1 | 1 | autoimmune disease of the blood |
| autoimmune pancreatitis | DTREE_00000110 | 3 | 1 | autoimmune disease of the gastrointestinal tract |
| dermatomyositis | DOID:10223 | 4 | 4 | autoimmune disease of the skin and connective tissue |
| juvenile ankylosing spondylitis | DTREE_00000040 | 4 | 4 | autoimmune disease of the musculoskeletal system |
| thyrotoxic exophthalmos | DOID:12362 | 4 | 1 | autoimmune disease of the eyes , ears, nose , and throat |
| hemolytic-uremic syndrome | DOID:12554 | 5 | 2 | autoimmune disease of the urogenital tract |
| lichen planus | DOID:9201 | 5 | 2 | autoimmune disease of the skin and connective tissue |
| vasculitis | DOID:865 | 6 | 1 | autoimmune disease of the skin and connective tissue |
| autoimmune polyendocrine syndrome type 1 | DOID:0050167 | 7 | 3 | autoimmune disease of the endocrine system |
| Lambert-Eaton myasthenic syndrome | DOID:0050214 | 8 | 2 | autoimmune disease of the musculoskeletal system |
| psoriatic arthritis | DOID:9008 | 8 | 4 | autoimmune disease of the musculoskeletal system |
| thrombotic thrombocytopenic purpura | DOID:10772 | 8 | 1 | autoimmune disease of the blood |
| autoimmune optic neuritis | DTREE_00000057 | 9 | 3 | autoimmune disease of the eyes , ears, nose , and throat |
| cicatricial pemphigoid | DOID:11655 | 9 | 2 | autoimmune disease of the skin and connective tissue |
| encephalomyelitis | DOID:640 | 10 | 1 | autoimmune disease of the central nervous system |
| relapsing polychondritis | DOID:2556 | 10 | 1 | autoimmune disease of the musculoskeletal system |
| acquired epidermolysis bullosa | DOID:4313 | 11 | 1 | autoimmune disease of the skin and connective tissue |
| autoimmune gastritis | DTREE_00000066 | 11 | 2 | autoimmune disease of the gastrointestinal tract |
| Guillain-Barre syndrome | DOID:12842 | 11 | 5 | autoimmune disease of the central nervous system |
| paraneoplastic polyneuropathy | DOID:8681 | 12 | 3 | autoimmune disease of the central nervous system |
| ulcerative colitis | DOID:8577 | 12 | 10 | autoimmune disease of the gastrointestinal tract |
| non-insulin-dependent diabetes mellitus | DOID:9352 | 16 | 6 | autoimmune disease of the endocrine system |
| alopecia areata | DOID:986 | 17 | 6 | autoimmune disease of the skin and connective tissue |
| stiff-person syndrome | DOID:13366 | 23 | 4 | autoimmune disease of the musculoskeletal system |
| Crohn's disease | DOID:8778 | 25 | 10 | autoimmune disease of the gastrointestinal tract |
| cryoglobulinemia | DOID:2917 | 26 | 4 | autoimmune disease of the blood |
| demyelinating polyneuropathy | DOID:5214 | 27 | 5 | autoimmune disease of the peripheral nervous system |
| autoimmune polyendocrine syndrome | DOID:14040 | 30 | 4 | autoimmune disease of the endocrine system |
| Vogt-Koyanagi-Harada | DOID:12297 | 33 | 4 | autoimmune disease of the skin and connective tissue |
| sclerosing cholangitis | DOID:14268 | 36 | 7 | autoimmune disease of the gastrointestinal tract |
| pemphigus gestationis | DOID:14482 | 40 | 1 | autoimmune disease of the skin and connective tissue |
| vitiligo | DOID:12306 | 41 | 6 | autoimmune disease of the skin and connective tissue |
| antiphospholipid syndrome | DOID:2988 | 42 | 6 | autoimmune disease of the blood |
| autoimmune haemolytic anaemia | DOID:718 | 44 | 3 | autoimmune disease of the blood |
| Addison's disease | DOID:13774 | 56 | 2 | autoimmune disease of the endocrine system |
| autoimmune vasculitis | DTREE_00000101 | 62 | 9 | autoimmune disease of the cardiovascular system |
| Goodpasture's syndrome | DOID:9808 | 62 | 11 | autoimmune disease of the urogenital tract |
| peripheral neuropathy | DOID:0060053 | 64 | 1 | autoimmune disease of the peripheral nervous system |
| ankylosing spondylitis | DOID:7147 | 65 | 36 | autoimmune disease of the musculoskeletal system |
| prediabetes syndrome | DOID:11716 | 65 | 13 | autoimmune disease of the endocrine system |
| bullous pemphigoid | DOID:8506 | 70 | 3 | autoimmune disease of the skin and connective tissue |
| neuromyelitis optica | DOID:8869 | 74 | 13 | autoimmune disease of the central nervous system |
| mixed connective tissue disease | DOID:3492 | 76 | 20 | autoimmune disease of the skin and connective tissue |
| rheumatic myocarditis | DOID:8481 | 77 | 6 | autoimmune disease of the musculoskeletal system |
| autoimmune uveitis | DTREE_00000058 | 81 | 12 | autoimmune disease of the eyes , ears, nose , and throat |
| autoimmune atherosclerosis | DTREE_00000060 | 82 | 2 | autoimmune disease of the cardiovascular system |
| reactive arthritis | DOID:6196 | 83 | 36 | autoimmune disease of the musculoskeletal system |
| autoimmune glomerulonephritis | DTREE_00000063 | 90 | 4 | autoimmune disease of the urogenital tract |
| psoriasis | DOID:8893 | 99 | 24 | autoimmune disease of the skin and connective tissue |
| Wegener's granulomatosis | DOID:12132 | 100 | 4 | autoimmune disease of the urogenital tract |
| immune thrombocytopenic purpura | DOID:8924 | 108 | 10 | autoimmune disease of the blood |
| juvenile rheumatoid arthritis | DOID:676 | 141 | 31 | autoimmune disease of the musculoskeletal system |
| primary biliary cirrhosis | DOID:12236 | 160 | 34 | autoimmune disease of the urogenital tract |
| pemphigus | DOID:9182 | 165 | 10 | autoimmune disease of the skin and connective tissue |
| autoimmune hepatitis | DOID:2048 | 218 | 16 | autoimmune disease of the gastrointestinal tract |
| Sjogrens syndrome | DOID:12894 | 250 | 39 | autoimmune disease of the exocrine system |
| autoimmune thyroiditis | DOID:7188 | 252 | 77 | autoimmune disease of the endocrine system |
| myasthenia gravis | DOID:437 | 303 | 16 | autoimmune disease of the musculoskeletal system |
| Grave's disease | DOID:12361 | 413 | 76 | autoimmune disease of the endocrine system |
| systemic scleroderma | DOID:418 | 446 | 57 | systemic autoimmune disease |
| Behcet syndrome | DOID:13241 | 646 | 547 | autoimmune disease of the eyes , ears, nose , and throat |
| insulin-dependent diabetes mellitus | DOID:9744 | 787 | 72 | autoimmune disease of the endocrine system |
| systemic lupus erythematosus | DOID:9074 | 909 | 128 | systemic autoimmune disease |
| celiac disease | DOID:10608 | 972 | 20 | autoimmune disease of the gastrointestinal tract |
| rheumatoid arthritis | DOID:7148 | 2049 | 416 | autoimmune disease of the musculoskeletal system |
| multiple sclerosis | DOID:2377 | 2187 | 473 | autoimmune disease of the central nervous system |
